# Lipidomics of homeoviscous adaptation to low temperatures in *Staphylococcus aureus* utilizing exogenous straight-chain unsaturated fatty acids over biosynthesized endogenous branched-chain fatty acids

**DOI:** 10.1101/2024.02.02.578686

**Authors:** Shannon C. Barbarek, Ritika Shah, Sharanya Paul, Gloria Alvarado, Keerthi Appala, Emma C. Henderson, Evan T. Strandquist, Antje Pokorny, Vineet K. Singh, Craig Gatto, Jan-Ulrik Dahl, Kelly M. Hines, Brian J. Wilkinson

## Abstract

It is well established that *Staphylococcus aureus* can incorporate exogenous straight-chain unsaturated fatty acids (SCUFAs) into membrane phospho- and glyco-lipids from various sources in supplemented culture media, and when growing *in vivo* in an infection. Given the enhancement of membrane fluidity when oleic acid (C18:1Δ9) is incorporated into lipids, we were prompted to examine the effect of medium supplementation with C18:1Δ9 on growth at low temperatures. C18:1Δ9 supported the growth of a cold-sensitive, branched-chain fatty acid (BCFA)-deficient mutant at 12°C. Interestingly, we found similar results in the BCFA-sufficient parental strain. We show that incorporation of C18:1Δ9 and its elongation product C20:1Δ9 into membrane lipids was required for growth stimulation and relied on a functional FakAB incorporation system. Lipidomics analysis of the phosphatidylglycerol (PG) and diglycosyldiacylglycerol (DGDG) lipid classes revealed major impacts of C18:1Δ9 and temperature on lipid species. Growth at 12°C in the presence of C18:1Δ9 also led to increased production of the carotenoid pigment staphyloxanthin; however, this was not an obligatory requirement for cold adaptation. Enhancement of growth by C18:1Δ9 is an example of homeoviscous adaptation to low temperatures utilizing an exogenous fatty acid. This may be significant in the growth of *S. aureus* at low temperatures in foods that commonly contain C18:1Δ9 and other SCUFAs in various forms.

**IMPORTANCE:** We show that *S. aureus* can use its known ability to incorporate exogenous fatty acids to enhance its growth at low temperatures. Individual species of phosphatidylglycerols and diglycosyldiacylglycerol bearing one or two degrees of unsaturation derived from incorporation of C18:1Δ9 at 12°C are described for the first time. In addition, enhanced production of the carotenoid staphyloxanthin occurs at low temperatures. The studies describe a biochemical reality underlying in membrane biophysics. This is an example of homeoviscous adaptation to low temperatures utilizing exogenous fatty acids over the regulation of the biosynthesis of endogenous fatty acids. The studies have likely relevance to food safety in that unsaturated fatty acids may enhance growth of *S. aureus* in the food environment.

## INTRODUCTION

The gram-positive bacterium *Staphylococcus aureus* is a versatile pathogen of humans and other animals that is capable of infecting various organs and tissues. *S. aureus* infections are often difficult to treat due to the accumulation of resistance to multiple antimicrobial agents. Additionally, foodborne intoxications are caused by enterotoxin-producing *S. aureus* strains in food resulting in toxin ingestion and subsequent disease symptoms (1, 2). Although *S. aureus* is typically thought of as a host-associated bacterium that is only subjected to small variations in temperature (3), the organism is also present in the environment (4), and in food (1, 2), where it is exposed to a much wider range of temperatures. In fact, *S. aureus* can grow from about 7 to 48°C (5).

To cope with such a wide range of temperatures, bacteria must maintain appropriate membrane fluidity (viscosity) to ensure continued cellular functions. The fatty acids of membrane glycerolipids are the major determinants of membrane fluidity and their biosynthesis is strictly regulated to provide a constant membrane fluidity when it is measured at the growth temperature, a process known as homeoviscous adaptation (6). Fatty acid shortening and increased biosynthesis of straight-chain unsaturated fatty acids (SCUFAs) occur in response to lower growth temperatures in *Escherichia coli* (7). In bacteria that contain branched-chain fatty acids (BCFAs) and straight-chain fatty acids (SCFAs), such as is typical of various gram-positive bacteria including *S. aureus*, fatty acid shortening and branching switching from iso to anteiso forms are the major adaptations to lower growth temperatures (8, 9).

The fatty acids of *S. aureus* are a mixture of BCFAs and SCFAs and the organism is unable to biosynthesize SCUFAs (10). The bacterium can incorporate exogenous SCUFAs and SCFAs as free fatty acids (11) (, from triglycerides and cholesteryl esters, complex biological materials including serum, and when growing in an infection (10, 12–16). Incorporation occurs via the FakAB system (11) and is thought to represent a savings of carbon and energy.

Given that *S. aureus* can incorporate SCUFAs such as C18:1Δ9 into its lipids, we were prompted to ask whether incorporation of this fatty acid into the membrane phospho- and glyco-lipids could play a role in the low-temperature adaptation of the organism. SCUFAs increase membrane fluidity more than BCFAs due to the geometry of the molecule with a *cis* double bond mid-chain, which thus packs less closely in the bilayer (17). We first examined this in a cold-sensitive strain deficient in BCFAs due to a transposon insertion in the branched-chain α-keto acid dehydrogenase (*bkd*) operon (18). Defective low-temperature growth of the BCFA-deficient strain was indeed stimulated by supplementation of medium with C18:1Δ9. However, C18:1Δ9 also markedly stimulated the growth of the parental strains with an intact *bkd* operon at low temperatures. C18:1Δ9 and its elongation product C20:1Δ9 were incorporated into lipids to significant amounts at all temperatures studied. In addition, the production of another membrane lipid component, the orange carotenoid staphyloxanthin, was induced by C18:1Δ9 at 12°C. SCUFAs are widely distributed in plant and animal material and may promote the growth of *S. aureus* at low temperatures in the food environment. These findings are an example of homeoviscous adaptation to low temperatures relying on exogenous fatty acids over the modulation of the biosynthesis of endogenous fatty acids, a phenomenon that does not appear to have been extensively considered in the literature.

## RESULTS

### C18:1Δ9 stimulates the growth of BCFA-deficient strain JE2Δ*lpd* at low temperatures

Strains with transposon insertions in the *bkd* operon are deficient in BCFAs (e.g., 35.4% BCFA versus 63.5% in the corresponding parental strain), and show increasingly diminished growth as the culture temperature is lowered (18). Two-methyl butyrate is a precursor of anteiso C15:0 and C17:0 fatty acids, which are particularly important in low-temperature growth (18). This compound stimulates the growth of BCFA-deficient mutants due to the restoration of the biosynthesis of anteiso fatty acids. For these reasons, we first studied the impact of C18:1Δ9 on the growth of this mutant at various temperatures (**Fig. 1**). C18:1Δ9 delayed growth and final optical densities were lower at 37°C (**Fig. 1a**) and 2-methyl butyrate had little impact on the growth of strain JE2Δ*lpd* at 37°C (**Fig. 1a**). In contrast, growth of JE2Δ*lpd* was much slower at 20°C and was markedly improved by C18:1Δ9 but not 2-methyl butyrate (**Fig. 1b**). At 12°C, growth of the BCFA-deficient strain was extremely slow but was promoted markedly by the presence of C18:1Δ9 (**Fig. 1c**). Thus, our data suggest that C18:1Δ9 was able to substitute for BCFAs at low temperatures and input sufficient fluidity to the membrane to promote growth. Fatty acid anteiso C15:0 added to the culture medium was inhibitory at all temperatures tested (data not shown), which is in agreement with the findings of Parsons et al. (11).

**Figure 1.**
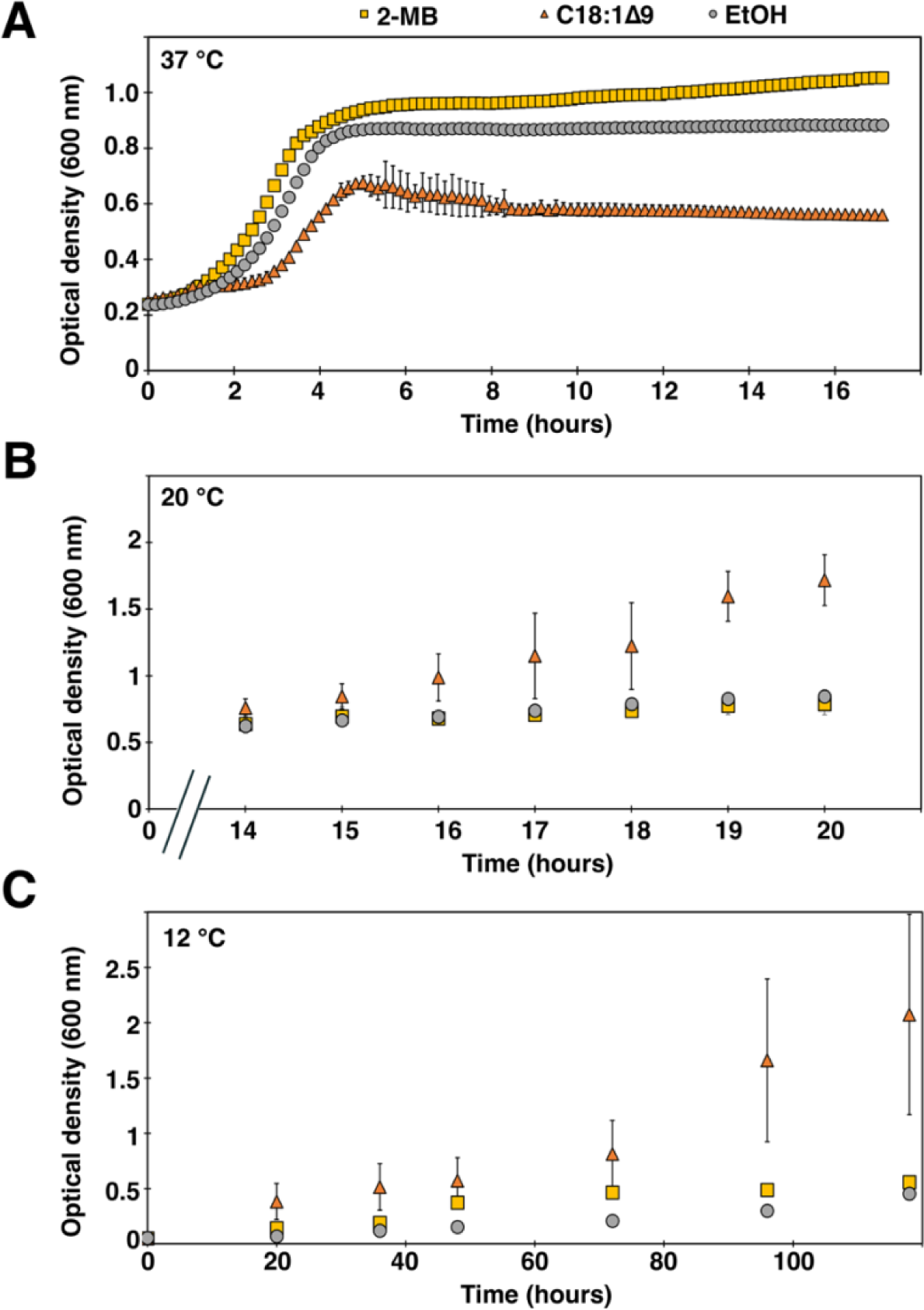
C18:1Δ9 stimulates growth at lower temperatures. Growth curve of JE2Δ*lpd* grown in the presence and absence of 100 μM 2-methyl butyrate and C18:1Δ9 at **(A)** 37 °C, **(B)** 20 °C, and **(C)** 12 °C, respectively. OD 600 nm was determined at the indicated time points (n = 3, ± S.D.).

### C18:1Δ9 stimulates the growth of the BCFA-sufficient parental strain JE2 at 12°C in a FakA-dependent manner

Given the stimulation of the growth of the BCFA-deficient strain at low temperatures, it was of interest to investigate this in the BCFA-sufficient parental strain that is known to incorporate C18:1Δ9 at 37°C (10). The addition of 70 µM C18:1Δ9 to cultures at 37°C delayed the growth of the culture by about two hours, and delayed growth to a small extent in cultures at 20°C (data not shown). However, somewhat to our surprise, while growth at 12°C was very poor in TSB alone, supplementation of medium with C18:1Δ9 supported marked growth of the culture (**Fig. 2**) suggesting that the growth promoting effect of C18:1Δ9 also occurs in the presence of normal levels of BCFA at low temperatures. The stimulation of growth at 12°C was dependent upon an intact FakA system because the growth of the *fakA*::Tn mutant was not stimulated by C18:1Δ9. This indicates that C18:1Δ9 and its elongation products must be incorporated into membrane phospho- and glyco-lipids to impart sufficient fluidity to the membrane enabling growth at 12°C.

**Figure 2.**
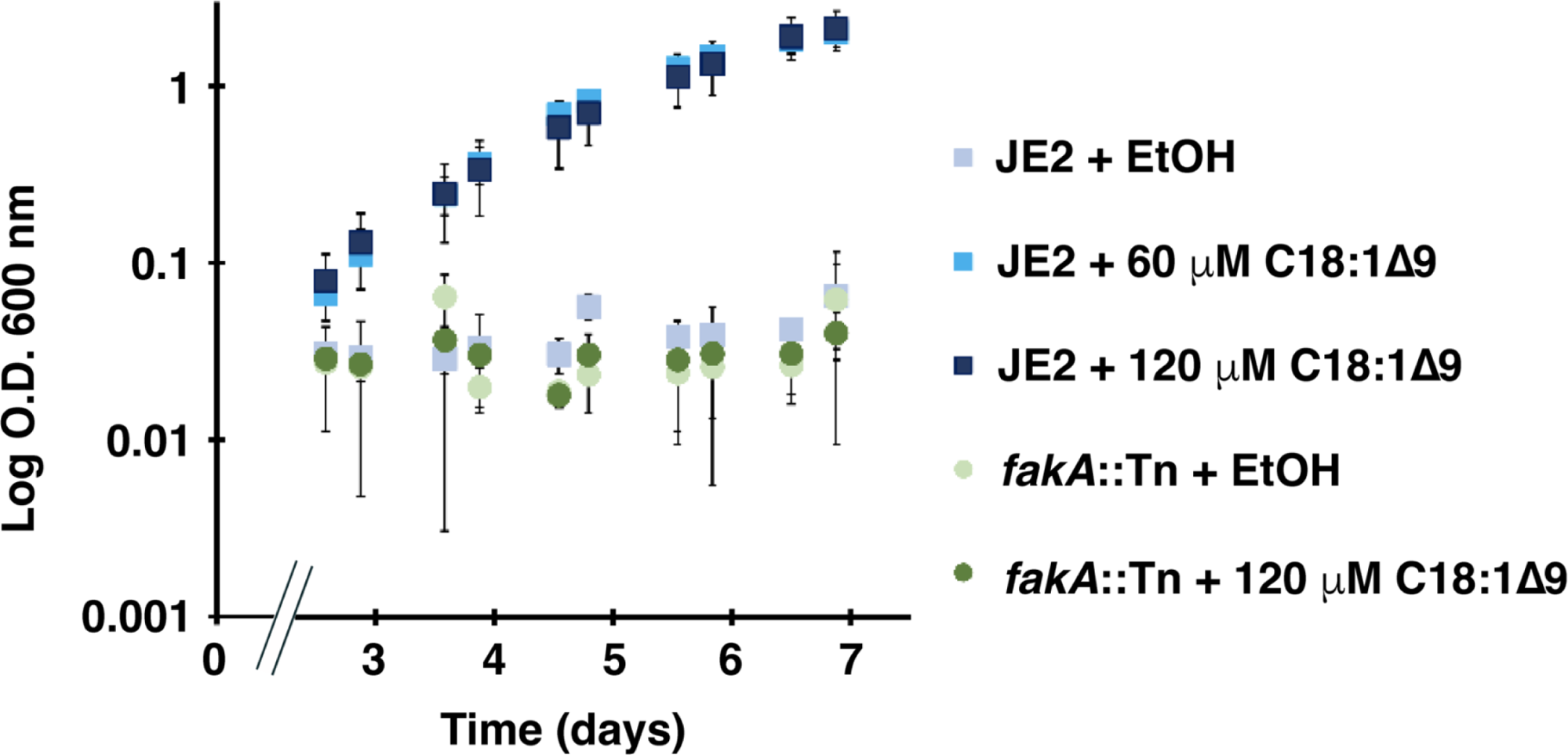
C18:1Δ9 promotes growth in a *fakA*-dependent manner. Growth curve of JE2 and *fakA*::Tn grown in the presence and absence of the indicated concentrations of C18:1Δ9. OD 600 nm was determined at the indicated time points (n = 3, ± S.D.).

### C18:1Δ9 is incorporated into cellular lipids in a FakA-dependent manner during growth at all temperatures

We first analyzed the major classes of fatty acids in JE2Δlpd grown at 12°C in the absence and presence of C18:1Δ9 and 2-methylbutyrate. Cells grown at 12°C had about 35% SCFAs and 65% BCFAs (considerable amounts of culture needed to be harvested because growth was very poor at this temperature). When cells were grown in the presence of 70 μM C18:1Δ9 SCUFAs made up 34% of total fatty acids. Inclusion of 2-methylbutyrate in the medium increased anteiso BCFAs from 27% in its absence to 83% in its presence. The SCUFAs were composed of C18:1Δ9 and its elongation product C20:1Δ9 (data not shown).

The compositions of the major classes of fatty acids of strain JE2 grown in TSB-supplemented with ethanol (control) or 70 μM C18:1Δ9 are shown in **Table 1**. These cells were composed of about 28% SCFAs and 62% BCFAs. SCUFAs were not present in cells grown in unsupplemented TSB. However, when grown in the presence of C18:1Δ9, SCUFAs made up 58.7% of the total fatty acid profile at 12°C (**Table 1**). At this temperature, anteiso fatty acids diminished from 49.5% in cultures grown without C18:1Δ9 to 19.6% in cells from C18:1Δ9-supplemented cultures. SCFAs decreased from 27.8% to 13.3% under these conditions. Similarly, growth at 20 °C resulted in incorporation of 71.2% SCUFAs in cells that were grown in the presence of C18:1Δ9. SCUFAs, which made up 29.5% of the total fatty acids in cells grown at 37°C in the presence of C18:1Δ9, were composed of 9.4% C18:1Δ9 and 19.6% of its elongation product C20:1Δ9. At 12°C in C18:1Δ9-supplemented cultures, C18:1Δ9 was 29.7% of SCUFAs and C20:1Δ9 was 28.9%, whereas at 20 °C, C18:1Δ9 was predominantly incorporated (**Table 1**). Hence, there was a tendency towards the shorter chain unsaturated fatty acid at low temperatures. C18:1Δ9 was incorporated to a negligible extent by the *fakA* mutant at 37°C, thus verifying that the FakA system is required for C18:1Δ9 incorporation into membrane lipids.

**TABLE 1.** Major fatty acid classes and individual fatty acids in strains JE2 and JE2Δ*fakA* grown in the presence and absence of C18:1Δ9 at different temperatures.

| Fatty Acid Classes and Major Fatty Acids | % Total Fatty Acid Composition<sup>a</sup> |  |  |  |  |  |  |
| --- | --- | --- | --- | --- | --- | --- | --- |
|  | Strains and Conditions |  |  |  |  |  |  |
| | JE2<br>12°C | JE2<br>12°C +<br>C18:1Δ9 | JE2<br>20°C | JE2<br>20°C +<br>C18:1Δ9 | JE2<br>37°C | JE2<br>37°C +<br>C18:1Δ9 | JE2Δ*fakA*<br>37°C +<br>C18:1Δ9 |
| **Even SCFAs** | **27.8 ± 0.8** | **13.3 ± 2.4** | **37.2 ± 0.2** | **12.8 ± 0.3** | **40.4 +/- 1.7** | **31.6 +/- 2.2** | **57.7 +/- 0.2** |
| C18:0 | 7.5 ± 0.1 | 2.3 ± 0.8 | 13.3 ± 0.1 | 3.1 ± 0.1 | 16.7 +/- 0.0 | 6.2 +/- 0.4 | 6.6 +/- 0.1 |
| C20:0 | 15.2 ± 1.2 | 6.6 ± 2.9 | 17.7 ± 0.0 | 6.3 ± 0.0 | 18.4 +/- 2.2 | 19.0 +/- 2.7 | 41.9 +/- 0.1 |
| **Iso odd BCFA**s | **12.5 ± 0.4** | **6.3 ± 1.3** | **18.9 ± 0.2** | **5.4 ± 0.1** | **16.0 +/- 0.6** | **12.1 +/- 0.1** | **11.3 +/- 0.0** |
| Iso C15:0 | 8.5 ± 0.4 | 4.9 ± 0.3 | 13.7 ± 0.3 | 5.0 ± 0.5 | 11.3 +/- 0.8 | 10.0 +/- 0.1 | 8.7 +/- 0.0 |
| **Anteiso odd BCFA**s | **49.5 ± 1.0** | **19.6 ± 0.8** | **34.3 ± 0.0** | **9.1 ± 0.8** | **36.0 +/- 1.1** | **22.9 +/- 0.2** | **20.7 +/- 0.0** |
| Anteiso C15:0 | 38.9 ± 1.3 | 18.1 ± 0.4 | 30.8 ± 0.2 | 8.9 ± 0.8 | 30.4 +/- 1.4 | 21.1 +/- 0.1 | 17.6 +/- 0.0 |
| **SCUFAs** | **N.D.** | **58.7 ± 1.8** | **N.D.** | **71.2 ± 1.0** | **N.D.** | **29.5 +/- 2.4** | **1.6 +/- 0.0** |
| C18:1Δ9 | N.D. | 29.7 ± 2.7 | N.D. | 62.3 ± 2.6 | N.D. | 9.4 +/- 1.1 | 1.6 +/- 0.0 |
| C20:1Δ9 | N.D. | 28.9 ± 3.8 | N.D. | 8.9 ± 1.5 | N.D. | 19.6 +/- 1.4 | N.D. |

<sup>a</sup> The percent total fatty acids in each major class are shown in bold and the major fatty acids in each class are shown.
<sup>b</sup> N.D. not detected

### Lipidomic analyses

C18:1Δ9 incorporation was detected in both the phosphatidylglycerol (PG) and diglycosyldiacylglycerol (DGDG) lipid classes for all three temperatures investigated (**Fig. 3 - 4**). The incorporation of C18:1Δ9 tended to suppress the levels of PGs and DGDGs with endogenous saturated fatty acids relative to the control conditions (**Fig. 3A and Fig. 4A**). The predominant species of PGs and DGDGs formed upon C18:1Δ9 incorporation had an odd-carbon fatty acyl tail in addition to the C18:1 or C20:1 fatty acid (**Fig. 3B and Fig. 4B**). Both PG 33:1 and DGDG 33:1 resulted from the pairing of C18:1Δ9 with C15:0 fatty acid, whereas PG 35:1 and DGDG 35:1 were formed from elongated C18:1Δ9 (C20:1) and C15:0. Bacteria grown with C18:1Δ9 also contained PG and DGDG 31:1. Both the 31:1 and 33:1 lipids show an inverse relationship between their abundance and the growth temperature, with levels being highest in the at 12°C cultures and lowest at 37°C. Lipids containing an even number of total acyl carbons and one degree of unsaturation (i.e., 32:1, 34:1, 36:1) were also detected, but tended to decrease as the growth temperature decreased. On the other hand, bacteria grown at 12°C in the presence of C18:1Δ9 formed substantially higher amounts of PGs and DGDGs containing two C18:1Δ9-derived fatty acids (**Fig. 3C,D and Fig. 4C**). PG 36:2 was over 9-times higher in the 12°C cultures than either the 20°C or 37°C cultures, while PG 38:2 was 3.5- and 4.7-times higher at 12°C than 20°C or 37°C, respectively. Although these overall trends were consistent between PG and DGDG species, the relative differences between the three growth temperatures were much larger for the DGDG species. For example, DGDG 36:2 was 14- and 30-times higher in bacteria grown at 12°C than 20°C or 37°C, respectively.

**Figure 3.**
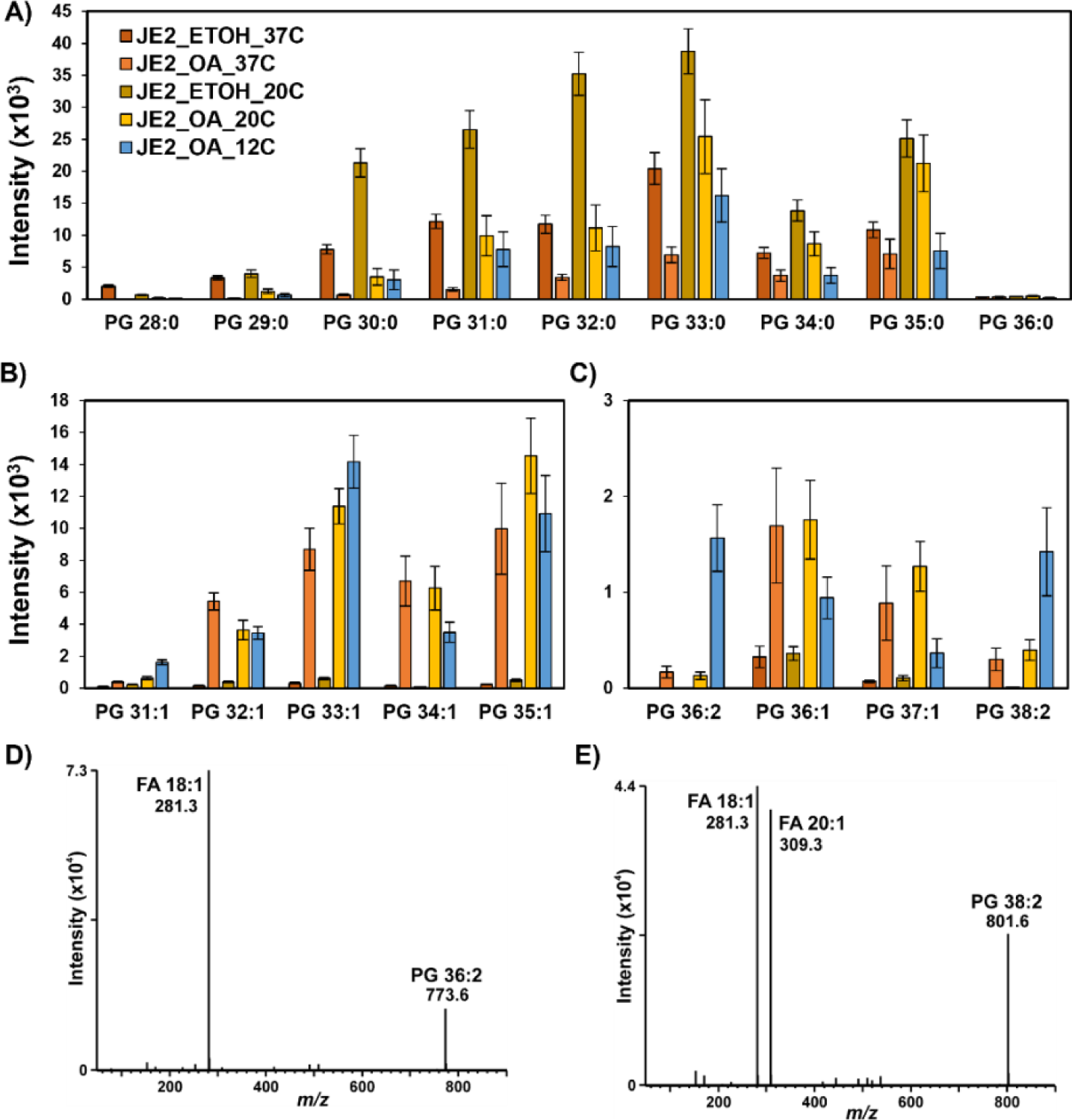
Lipidomics analysis of phosphatidylglycerol (PG) lipids in JE2 grown at 12, 20, and 37°C with and without oleic acid. A) PGs containing saturated fatty acids; B) high-abundance PGs containing monounsaturated fatty acids; C) low abundance PGs containing one or two monounsaturated fatty acids; D) Negative mode fragmentation spectra of PG 36:2, m/z 773.6, showing the presence of only FA 18:1 acyl tails; E) Negative mode fragmentation spectra of PG 38:2, m/z 801.6, showing the presence of FA 18:1 and FA 20:1 acyl tails.

**Figure 4.**
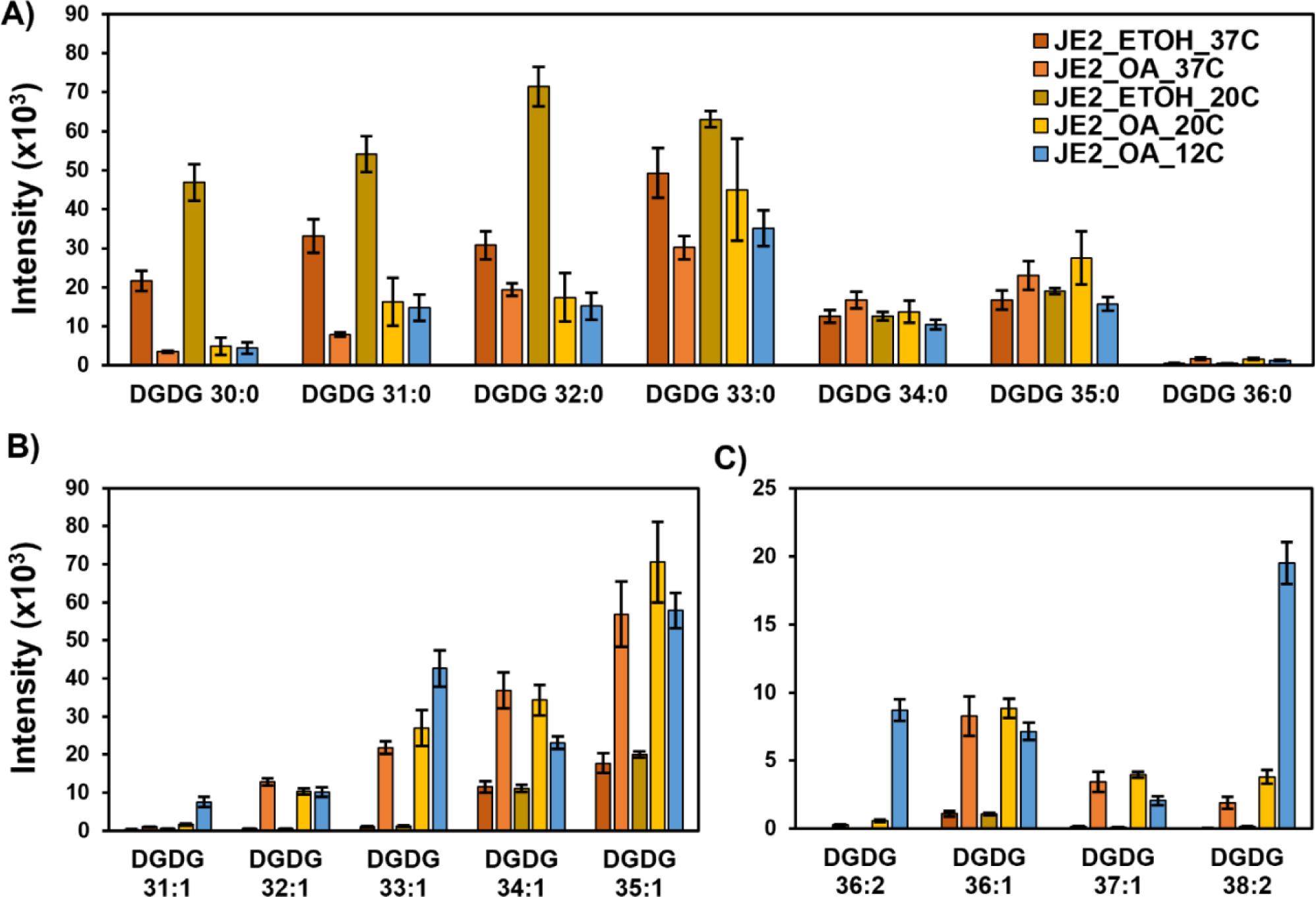
Lipidomics analysis of diglycosyldiacylglycerol (DGDG) lipids in JE2 grown at 12, 20, and 37°C with and without oleic acid. A) DGDGs containing saturated fatty acids; B) high-abundance DGDGs containing monounsaturated fatty acids; C) low abundance DGDGs containing one or two monounsaturated fatty acids.

In addition to the incorporation of C18:1Δ9 into PGs and DGDGs, temperature dependent effects were observed in the abundance of 10-hydroxystearic acid that is produced by the activity of oleate hydratase (Ohy) on C18:1Δ9. Oleate hydratase adds water to *cis*-9 double bonds, provides protection against palmitoleic acid (C16:1Δ9), and is considered to be a *S. aureus* virulence factor (19). Levels of 10-hydroxystearic acid were more than 6-fold lower in bacteria grown with C18:1Δ9 at 12°C and 20°C than at 37°C (**Fig. 5**). These data suggest that Ohy is less active at low temperatures, thereby promoting greater incorporation of C18:1Δ9 into membrane lipids at low temperatures.

**Figure 5.**
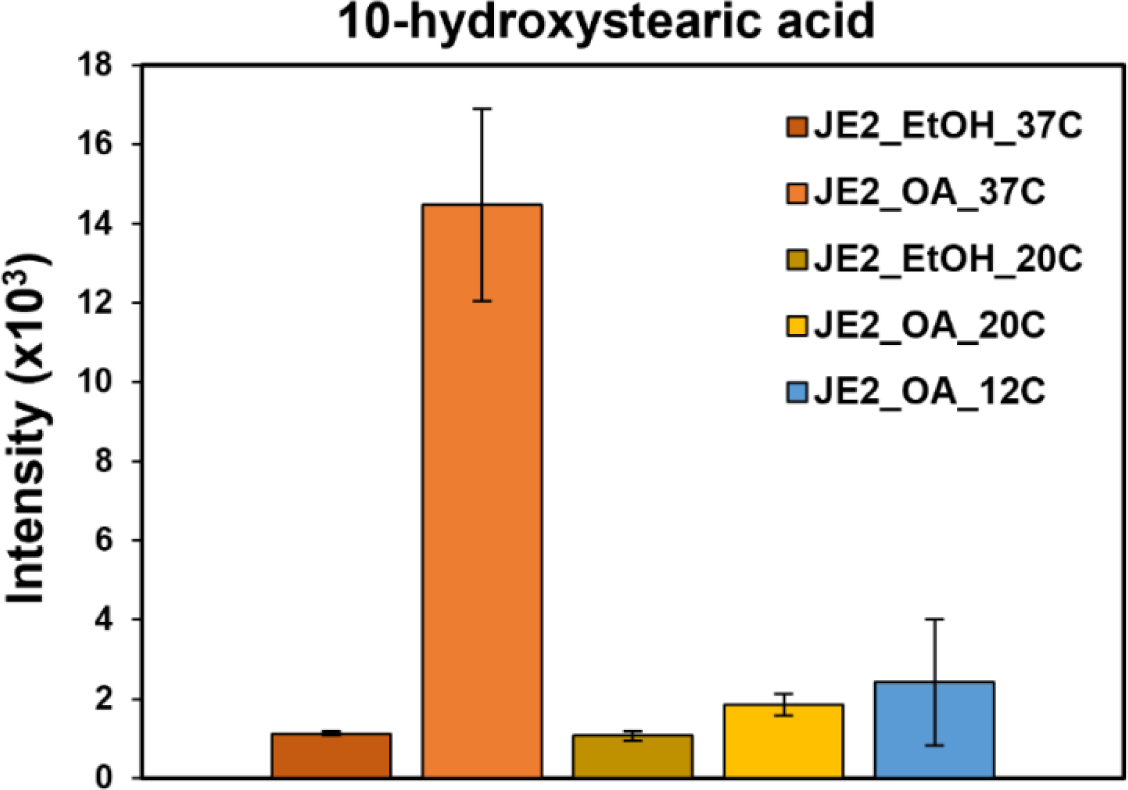
Abundance of 10-hydroxystearic acid, produced from oleic acid, in JE grown at 12, 20, and 37°C with and without oleic acid.

### C18:1Δ9 stimulated the production of staphyloxanthin at 12°C

When cultures were harvested for the determination of total fatty acid compositions, we noted that cultures grown at 12°C in the presence of C18:1Δ9 had a deeper orange color compared to cells grown in either its presence or its absence at higher temperatures. The results of quantification of staphyloxanthin levels are shown in **Fig. 6**. Staphyloxanthin levels were about four-fold higher in cells grown at 12°C with C18:1Δ9 compared to cells grown in either its presence or absence at 37°C. Thus, an increase in staphyloxanthin content appears to be involved in cold temperature adaptation in *S. aureus.* Increased production of staphyloxanthin at 25°C compared to 37°C has previously been noted by Joyce and others (20). It was thus of interest to study the stimulation of low-temperature growth in a carotenoid-deficient mutant. C18:1Δ9 stimulated the growth of carotenoid-deficient mutant JE2Δ*crtM* and the complemented strain JE2Δ*crtMc* pCU_crt_*OPQMN* to a similar extent at 12°C (data not shown). Thus, although staphyloxanthin production is enhanced at low temperatures by C18:1Δ9, staphyloxanthin production is not an obligatory part of the response to low temperatures. The carotenoid-deficient strain and its complement showed a high proportion of SCUFAs similar to strain JE2 when grown in the presence of C18:1Δ9 at 12°C (data not shown).

**Figure 6.**
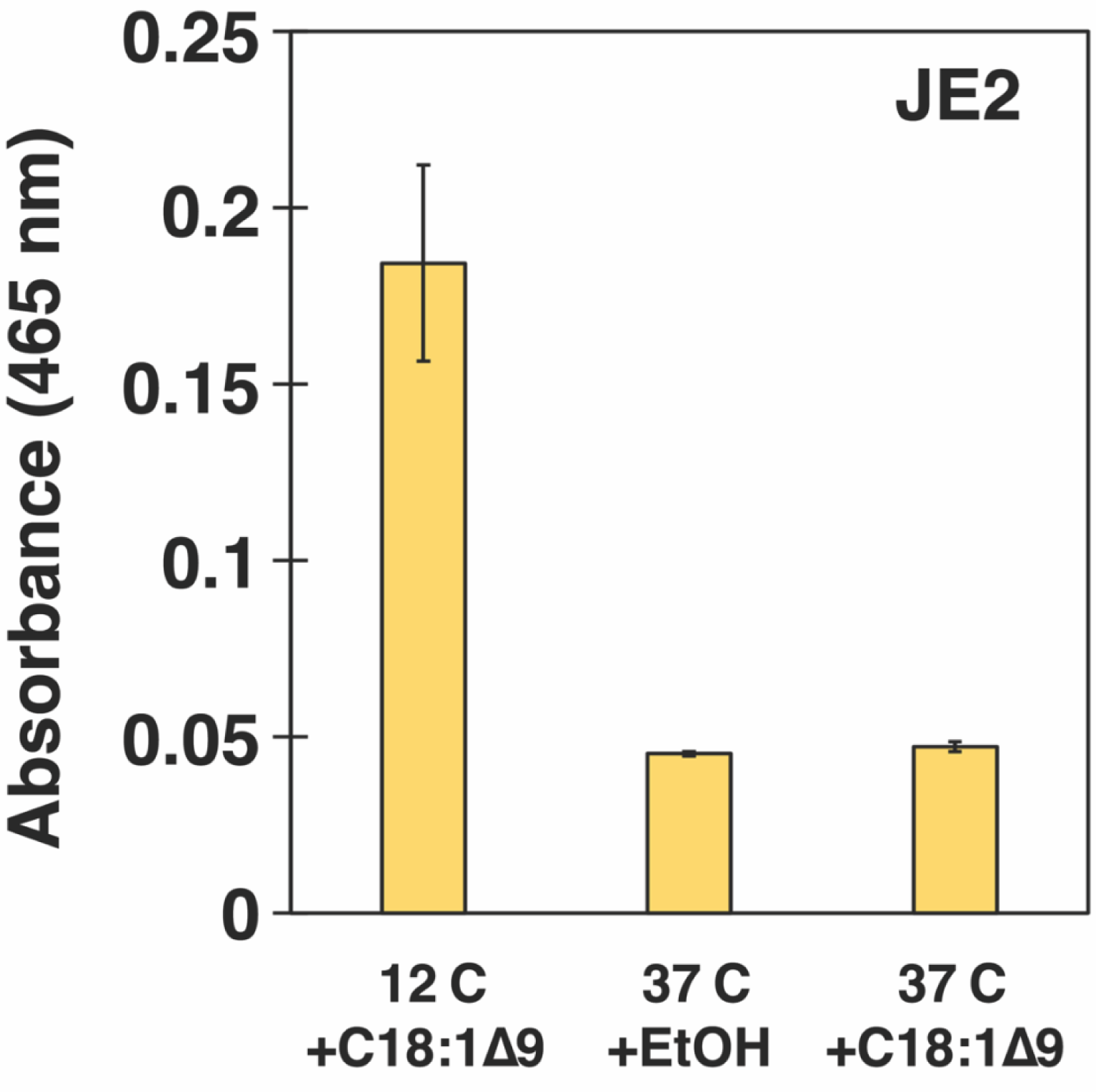
Staphyloxanthin content of *S. aureus* grown at different temperatures in the presence and absence of 100 μM C18:1Δ9. The indicated cultures were collected on day 5, washed twice in PBS, and normalized to 5 ml of an O.D._600 nm_ = 2. Staphyloxanthin was extracted with warm methanol (55°C) for 5 min and the O.D._465 nm_ of the supernatant was determined.

## DISCUSSION

BCFA-deficient mutants of *S. aureus* have lowered BCFAs and increased SCFAs and show increasingly impaired growth as the temperature lowers due to insufficient membrane fluidity (18). Growth at low temperatures can be rescued by feeding 2-methylbutyrate that acts as a precursor of anteiso fatty acids, in particularly fatty acid anteiso C15:0, that imparts fluidity to the membrane due to its lower degree of packing compared to SCFAs. We found that the SCUFA C18:1Δ9 was also able to promote growth of the BCFA-deficient mutant at low temperatures through its direct incorporation into membrane lipids.

It is well known that *S. aureus* incorporates exogenous SCUFAs into its phospho- and glyco-lipids from various sources(10, 12–16, 21). We found that C18:1Δ9 caused a minor delay in culture growth at 37°C and that substantial incorporation of SCUFAs into membrane lipids occurred at 37°C and 20°C. Growth of strain JE2 at 12°C was very slow, but C18:1Δ9 markedly stimulated growth. This growth stimulation was dependent on incorporation via the FakAB system because a FakA-deficient strain did not grow at 12°C in the presence of C18:1Δ9.

C18:1Δ9, owing to its kinked *cis*-double bond structure, disrupts membrane packing and increases membrane fluidity to a greater degree than the iso- and anteiso-BCFAs present in the *S. aureus* membrane under normal growth conditions in complex media (17). Indeed, phospholipids containing C18:1Δ9 demonstrate a melting transition around or below the freezing point of water (22, 23). C18:1Δ9 then clearly impacts the adaptation of *S. aureus* to lower temperatures, a phenomenon known as homeoviscous adaptation. The appearance of mono and double unsaturated species of PGs and DGDGs in the lipid profiles at low temperatures is a result of C18:1Δ9 incorporation and its elongation product by the FakAB system.

### Homeoviscous adaptation using an exogenous fatty acid over or in addition to the regulation of the biosynthesis of endogenous fatty acids

Sinensky (6) showed that bacteria have cytoplasmic membranes that show a constant fluidity when fluidity is measured at the growth temperature. This is achieved through the regulation of the biosynthesis of endogenous fatty acids. The strategies involved in achieving homeoviscous adaptation vary depending on whether the fatty acids of an organism are predominantly SCFAs and SCUFAs such as is found in *Escherichia coli* and other gram-negative bacteria, or contain significant amounts of BCFAs and SCFAs typical of many gram-positive bacteria (24, 25). In the case of SCFA-SCUFA-containing bacteria, SCFAs decrease and SCUFAs increase as temperatures decrease (25, 26). In the absence of such adaptation, the membrane fluidity would decrease, and the membrane would become too rigid for continued cellular function and growth could cease. In BCFA-containing bacteria such as *Listeria monocytogenes* and *Bacillus subtilis,* fatty acid shortening and branching switching to anteiso fatty acids are the main strategies in response to lower growth temperatures (8, 25). have provided evidence that in *E. coli* homeoviscous adaptation involves a temperature-responsive metabolic valve regulating flux between the pathways for biosynthesis of SCFAs and SCUFAs. This is regulation of the biosynthesis of endogenous fatty acids to achieve appropriate membrane fluidity. A temperature-responsive regulation of a transcription-based negative feedback loop is a second element in homeoviscous adaptation. The picture is less clear in BCFA-containing bacteria but the initial committed enzyme in fatty acid biosynthesis (FabH) in *L. monocytogenes,* shows increased preference for C5-CoA precursors of anteiso fatty acids at low temperatures (27).

### The incorporation of SCUFAs by *S. aureus* adds exogenous SCUFAs as a homeoviscosity strategy to the endogenous BCFA strategy

The membrane lipid changes in *S. aureus* in response to lower growth temperatures have not been studied extensively. Joyce et al. (20) showed significant increases in fatty acid anteiso C15:0 and the production of staphyloxanthin when cultures were shifted from 37 to 25°C. The importance of BCFAs to cold adaptation in *S. aureus* is demonstrated by the cold sensitivity of BCFA-deficient mutants (18). These authors also showed that BCFAs increased, particularly fatty acid anteiso C15:0, and SCFAs, particularly C18:0, decreased in cultures grown at lower temperatures. Similar changes are seen in the data presented here where BCFAs increase markedly in cells grown at 12°C, although growth is very slow. However, when C18:1Δ9 is available, it is utilized and incorporated into membrane lipids and permits much more vigorous growth at low temperatures than can be achieved by BCFAs alone. This is presumably due to the greater fluidity imparted by this fatty acid with its unsaturated double bond in the middle of the molecule compared to the branching at the end of the molecule of BCFAs. Incorporation of C18:1Δ9 into *L. monocytogenes* lipids from the free fatty acid, polysorbate 80, and food lipid extract has been shown by Flegler et al. (28) and Touche et al.(29).. The mechanisms involved where the incorporation of an exogenous SCUFA overrides the modulation of the biosynthesis of endogenous BCFAs remain to be determined.

### Staphyloxanthin-another player in the staphylococcal homeoviscous response?

Staphyloxanthin is a golden triterpenoid pigment that is produced by *S. aureus*. The production of staphyloxanthin is vexingly variable amongst different strains and under different growth conditions (30, 31). The evidence that staphyloxanthin plays a role in protection against oxidative stress is reasonably convincing (32, 33). The general consensus is that staphyloxanthin rigidifies membranes (34). However, the finding of increased staphyloxanthin with decreased growth temperatures as reported here and previously by Joyce et al. (20)is surprising at first sight. Nevertheless, there are well-established connections between staphyloxanthin and gene expression in *S. aureus* in response to low temperatures. Katzif and others (35) reported that cold-shock protein A null mutants were white in color due to lack of staphyloxanthin production. CspA is proposed to act through a mechanism dependent on the alternative sigma factor SigB and was shown to be required for the expression of the carotenoid biosynthesis operon and SigB. CspA is believed to be an RNA chaperone (36). Increased production of staphyloxanthin and other carotenoids has been reported in the coagulase-negative staphylococcal species *Staphylococcus xylosus* grown at 10°C versus cultures grown at 30°C (37).

### Biophysical implications of the changes in membrane biochemistry at 12 °C

Seel et al. (37) carried out an in-depth biophysical investigation on the role of staphyloxanthin in membrane fluidity in *Staphylococcus xylosus*. Measurement of generalized polarization with Laurdan GP and anisotropy with trimethylammonium diphenylhexatriene (TMA-DPH) demonstrated simultaneous increases in membrane order and membrane fluidity correlated with increased carotenoid production. Increased expression of *crt* genes in low-temperature cultures was also observed. It seems likely that staphyloxanthin is playing a similar role in membrane adaptations to low temperatures in *S. aureus*. Múnera-Jaramillo et al. (38) have shown that staphyloxanthin reduces the liquid-crystalline to gel phase transition in *S. aureus* model membrane systems without significantly broadening the transition. This is indicative of a freezing-point depression caused by staphyloxanthin preferentially partitioning into the liquid-crystalline phase. However, we found that a carotenoid-deficient mutant also showed stimulation of growth at low temperatures by C18:1Δ9. This observation strongly suggests that staphyloxanthin serves to augment cold adaptation if present but is not strictly required. Moreover, based on these observations, we postulate that any membrane component that either softens the gel phase or lowers the membrane liquid-crystalline to gel phase transition temperature will aid bacterial survival at low temperatures.

### Food safety and low temperature process considerations

C18:1Δ9 is the most abundant fatty acid in nature and is found in various oils and fats, meats, cheese, eggs, and other foods in the form of free C18:1Δ9 and esterified to glycerol and cholesterol in the form of triglycerides and cholesteryl esters respectively. *S. aureus* clearly encounters C18:1Δ9 in multiple different foods, and it may enhance the growth of the organism at low temperatures. Staphylococcal food poisoning is a foodborne intoxication requiring initial contamination of food by *S. aureus* and then growth of the organism and release of enterotoxin. Storing food in refrigerators at 5°C should prevent the growth of *S. aureus*, but many domestic refrigerators are operated at higher temperatures than this, and maintaining the integrity of the cold chain is essential for the prevention of the growth of the organism in foods (39). It would seem likely that C18:1Δ9 either free or esterified, and possibly other unsaturated fatty acids, could enhance the growth of *S. aureus* above 6°C in foods. Additionally, a significant number of foods are produced by low-temperature fermentation processes (40). Possibly these fermentation processes and other low-temperature industrial applications could be enhanced by C18:1Δ9 or other unsaturated fatty acids by promoting more efficient exogenous homeoviscous adaptation of membrane fluidity leading to enhanced growth. As an example of this, Tanet al.(41) showed that exogenous unsaturated fatty acids enhanced the growth of *Lactobacillus casei* at low temperatures in the ripening of cheddar cheese.

## MATERIALS AND METHODS

### Bacterial strains and growth conditions

*S. aureus* strains used in this study are shown in **Table 2**. They were grown in 50ml of Tryptic Soy Broth (Difco) in 250ml Erlenmeyer flasks with shaking (200 rpm) at the specified temperature. Inocula were 2% from a starter culture grown overnight at 37°C. Growth was measured at intervals by measuring the optical density at 600 nm. Stock solutions of C18:1Δ9 and 2-methyl butyrate were prepared in 95% (v/v) ethanol and ethanol alone was added to cultures as a control. At least three biological replicates were performed for growth experiments.

**TABLE 2.** List of *S. aureus* strains used in this study.

| Strain | Characteristics | Reference |
| --- | --- | --- |
| JE2 | Derived from community-acquired MRSA strain USA 300 LAC | Fey et al., 2013 |
| JE2Δ*fakA* | NR-46772 | Fey et al., 2013 |
| JE2Δ*lpd* | BCFA-deficient mutant | Singh et al., 2018 |
| JE2Δ*crtM* | Carotenoid deficient mutant | Braungardt, H., & Singh, V. K. 2019 |
| JE2Δ*crtM* +<br>pCUcrtOPQMN | *crtM* mutant of JE2 complemented with *crt* gene cluster in *trans* | Singh et al., 2018 |

### Analysis of fatty acid composition

Cells were harvested by centrifugation and washed in cold 0.9% (w/v) NaCl solution before fatty acid analysis. Fatty acid methyl ester analysis was performed as described by Sen and others (10) at Microbial ID Inc (Newark, DE) where the cells were saponified, methylated, and extracted. The methyl ester mixtures were separated using an Agilent 5890 dual-tower gas chromatograph and the fatty acyl chains were analyzed and identified by the MIDI microbial identification system (Sherlock 4.5 microbial identification system). The percentages of the different fatty acids reported in tables and figures are the means of the values from two or three independent experiments. Some minor fatty acids are not reported.

### Estimation of carotenoid content

Washed harvested cells were extracted with warm methanol (55°C) for 5 min and the OD_465_ of the supernatant was measured as described before (10, 42).

### Lipid Extraction

The lipid extraction was performed using a modified version of the Bligh & Dyer extraction method (43, 44). The pellets were washed twice with 2 mL of sterile water, during which a portion of the suspension was taken for optical density measurement at 600 nm. After the last wash, the pellet was resuspended in 0.5 mL of HPLC grade water, transferred to glass culture tubes, and sonicated in an ice bath for 30 minutes. Chilled extraction solvent [1:2 chloroform/methanol (v/v)] was added to the samples and vortexed sporadically for 5 min. Phase separation was induced by adding 0.5 mL of chloroform and 0.5 mL of water, followed by 1 min of vortexing. The samples were centrifuged at 2,000 × g at 4° C for 10 min. The bottom layer containing the lipids was collected into fresh glass tubes and vacuum-dried (Savant, ThermoScientific). The dry extracts were reconstituted in 0.5 mL of 1:1 chloroform/methanol and stored in a −80°C freezer until the day of analysis.

### Liquid-chromatography and mass spectrometric analysis

Lipid extract aliquots were dried using a vacuum concentrator before being reconstituted in mobile phase A (MPA) at dilutions of 25X or greater. The samples were separated chromatographically with a Waters Acquity Ultra Performance Liquid Chromatography (UPLC) bridged ethylene hybrid (BEH) hydrophilic interaction liquid chromatography (HILIC) column (2.1 X 100mm, 1.7µm) at a constant temperature of 40°C. MPA composition consisted of 95% acetonitrile and 5% water, with 10 mM ammonium acetate, while mobile phase B (MPB) comprised a mixture of 50% acetonitrile and 50% water, containing 10 mM ammonium acetate. A constant flow rate of 0.5 mL/min was maintained throughout the 7 min run time. The gradient was as follows: 100% MPB for 0-2 min, decreasing to 60% MPB for 3-4 minutes, hold at 60% MPB for 4-5 min, and then returning to 100% MPB for 5-7 minutes. The autosampler was maintained at 6°C throughout the analysis. An injection volume of 5 µL was used.

After chromatographic separation, the samples were directed into the electrospray spray ionization (ESI) source of the Waters Synapt XS traveling-wave ion mobility-mass spectrometer (TWIMS-MS). The following variables were used for both negative and positive mode ionizations: capillary voltage, ±2 kV; source temperature, 120°C; desolvation temperature, 450°C; desolvation gas flow, 700 L/hr; cone gas flow, 50 L/hr. Ion mobility separation was performed in nitrogen (90 mL/min flow) with traveling wave settings of 550 m/s and 40V. Leucine enkephalin was used as a standard mass calibrant to lock mass correct the samples. Data was collected over 50–1200 *m/z* with a scan time of 0.5 sec. Data-independent acquisition (DIA) MS/MS experiments were conducted where fragmentation occurred in the transfer region of the instrument using a 45-60 eV collision energy ramp.

### Data Analysis

The Waters .raw files were imported into Progenesis QI software (v3.0, Waters/Nonlinear Dynamics). Subsequently the data was lock-mass corrected and aligned with the quality control (QC) reference file. Peak picking was performed using built-in default parameters. The data was normalized using OD600 readings to take into consideration the growth variations of *S. aureus* replicates. PGs (as [M − H]^−^ adducts) were evaluated from the negative mode ionization whereas DGDGs (as [M + NH4]^+^ adducts) and LysylPGs (as [M + H]^+^ adducts) were analyzed from the positive ionization mode with accurate mass (< 10 ppm tolerance) using LipidPioneer; user-generated database (45).

## Acknowledgements

This work was supported by National Institute of Health grants 1R21AI13535 to BJW and CG, R15AI164585 and 1R03AI174033-01A1 to JUD, and R01AI173144 to KMH, VKS and BJW. We thank Patrick Ofori Tawiah for carefully proof-reading the manuscript.

